# Implications of the trade-offs between negative density-dependence and Allee effects for vector control

**DOI:** 10.64898/2026.03.07.710288

**Authors:** Andrea M. Kipingu, Samson S. Kiware, Daniel T. Haydon, Paul C. D. Johnson, Mafalda Viana

**Affiliations:** School of Biodiversity, One Health and Veterinary Medicine, University of Glasgow, Graham Kerr Building, Glasgow G12 8QQ, United Kingdom; Environmental Health & Ecological Sciences Department, Ifakara Health Institute, P.O. Box 78 373, Dar es Salaam, Tanzania; The Pan-African Mosquito Control Association, KEMRI Headquarters, Mbagathi Road, Nairobi, Nairobi 54840-00200, Kenya

**Keywords:** Anopheles mosquitoes, larvicides, negative density-dependence, Allee effects, vector control, simulation model and population dynamics

## Abstract

**Background:** Understanding population dynamics is fundamental to predicting species persistence and extinction, yet remains challenging due to the complex interplay between ecological, environmental and anthropogenic factors. Population dynamics are regulated by intrinsic factors such as negative density-dependence and Allee effects. While negative density-dependence is a well-understood process, Allee effects have received less attention but can have important implications for conservation and species management. For control of disease vectors, negative density-dependence and Allee effects can drive vectors to elimination or to rebound after control. For malaria mosquitoes, negative density-dependence at the larval stages is well known to limit population growth, but the implications of Allee effects and the trade-offs between negative density-dependence and Allee effects remain unknown. It is hypothesised that, depending on the vector control strategy, Allee effects could be triggered and push populations closer to extinction.

**Methods:** A stochastic state-structured population simulation model was developed, which followed a simplified mosquito life cycle. Negative density-dependence and Allee effects were included as parameters influencing larval survival and total fecundity, respectively. The aims were addressed by varying the strength of negative density-dependence and Allee effects parameters, independently and simultaneously, under different vector control scenarios: control, sustained and shorter-term interventions (here, immolating larvicides) that reduced the larval population.

**Results:** While in isolation, the strength of negative density-dependence and Allee effects did not have a long-term impact on the population dynamics, their combination accelerated population extinction as both the strength of negative density-dependence and Allee effects increased. As Allee effects act on small population sizes, sustained interventions were able to activate Allee effects, increasing the probability of population extinction, but short-term interventions can lead to populations rebounding, driven by negative density-dependence.

**Conclusion:** Understanding less-studied regulatory processes like Allee effects can support vector control by highlighting aspects of the vector’s life cycle that are either resilient or vulnerable to interventions. If present, Allee effects could potentially be harnessed to accelerate the elimination of vectors and diseases such as malaria.

## Background

As populations decline, alterations in their per capita growth rate are likely to occur [1]. Two ecological processes may underpin population changes: negative density-dependence (DD), which stabilises populations at high densities through mechanisms such as resource limitation, and Allee effects (AE), which act to further reduce populations at low densities due to factors including mate limitation [2,3]. Vector control, a widespread strategy against many of the most important mosquito-borne diseases worldwide, including malaria, dengue, Zika, Rift Valley and yellow fever [4,5], typically works by reducing the mosquito population size and density. Understanding how population regulatory processes such as DD and AE are triggered by vector control when populations decline can then provide insights into how these might impede or facilitate vector control. This is particularly relevant for malaria control, the mosquito-borne disease with the highest global burden on human health.

It has been shown that in settings with low mosquito densities, malaria transmission can persist at low levels, despite pressure from different vector interventions. For example, in Tanzania, a large-scale larvicidal control in urban Dar es Salaam (Dar) reduced the Anopheles gambiae mosquito population by 94.4%, but malaria persisted [8–10]. Similarly, in Manica District, Mozambique, despite 69.2% of households having at least one insecticide-treated net, malaria transmission persisted [11]. More widely, malaria mosquito populations between 2000 and 2015 declined due to vector interventions [12], but were difficult to knock down further or eliminate, allowing malaria to persist at low levels. This could indicate that vector control interventions become proportionately less effective at low population densities, as surviving individuals can have higher per capita reproduction or growth rates than those in higher density populations.

It is well known that mosquito populations exhibit DD at the larval stages [13], with the per capita growth rate fastest at low densities. Laboratory experiments with Anopheles gambiae mosquitoes showed that resource competition is the major source of DD [13]. With less space (e.g., small water bodies), increased competition leads to larval population decline, which in turn releases some space to accommodate more larvae. Consequently, an intervention might become proportionately less effective as a population declines because surviving individuals grow more rapidly at low density. However, a positive scenario is also possible, where a small population experiences AE, leading to reduced population growth as its size declines [14]. AEs result from factors including mate limitation, limited cooperation, inbreeding and predation, leading to a higher risk of stochastic extinction as populations decline [2,15] and may operate at low or high levels. At low levels of AE, the population growth rate reduces but remains positive. Conversely, at higher levels of AE, growth rate may become negative, leading to population extinction [16]. As such, both DD and AE are expected to be important in regulating small populations. Examples of AE in other systems are provided in Supporting information (SI).

Although there is currently no evidence from field or laboratory experiments regarding AE in malaria mosquitoes or other vectors, previous modelling work predicted that gene drive could be more effective in low-density populations with strong AEs when transgenic mosquitoes are released [17]. There is currently a lack of understanding of how DD and AE trade-offs in mosquito populations and how these impact vector control. For instance, malaria poses a huge public health challenge, with its burden affecting millions worldwide [18]. While malaria vector control has been highly effective at reducing malaria mosquito populations [19], complete elimination remains rare. Understanding how DD and AE trade-off to regulate malaria mosquito populations could enhance malaria control and elimination efforts, particularly when targeting areas with low population densities.

This study aimed to address this knowledge gap by developing a stochastic simulation model based on a stage-structured population model to address three research questions: (i) What are the roles of DD and AE in regulating Anopheles gambiae mosquito populations at low densities? (ii) How do DD and AE impact the efficacy of the vector control interventions? and (iii) What are the trade-offs between DD and AE in regulating mosquito populations?

## Methods

### Study design

Tanzania remains among the sub-Saharan African countries with a high malaria burden, ranking seventh globally in terms of cases (3.3%) and among the four countries responsible for just over half of all malaria deaths (4.3%) [19]. Following the larvicidal programme in Dar, mosquito densities and malaria prevalence declined, after which transmission has persisted at low levels [20,21]. For this study, settings with low mosquito population abundances were required, and Dar was among the few regions where such declines were observed. Consequently, it was sought to emulate a population in which it would be biologically plausible to detect AE. A stochastic stage-structured simulation model of Anopheles gambiae mosquito population in Dar was developed, and its response to a larvicidal intervention was assessed. The larviciding in Dar [9] targeted 15 of 73 wards across 3 municipalities (5 wards each) and was deployed in three different phases (time periods) for 193 weeks from early 2005 to late 2008 [9]. In phase one, only three wards received a larvicidal treatment; in the second phase, another 6 wards were added, and in the last phase, the intervention was scaled up to cover all 15 wards. The type and amount of larvicides applied and intervention durations are described elsewhere [8,9,22]. Daily rainfall and temperature data from 2005-2008 were obtained from the open-source repository Soil and Water Assessment Tool [23] and aggregated to weekly means to match the time unit in the model (SI). Since the rainfall and temperature data are not specific for each ward, it was assumed that the mean weekly rainfall and temperature in all fifteen wards from the three municipalities of urban Dar were identical.

The simulation model was validated against observed female adult mosquitoes collected during the larvicidal intervention in Dar. The validation focused on assessing whether the model reproduced the temporal dynamics of the observed data, rather than on matching the absolute magnitudes of the simulated and observed abundances. To facilitate assessment across wards where differences in abundance magnitudes were substantial, both observed and simulated abundances were normalised by dividing each abundance by its maximum value over study weeks, resulting in normalised values ranging between 0 and 1. Model fit was further evaluated by plotting simulated against observed abundances to assess their match relative to the 1:1 line, providing a visual measure of goodness of fit.

### Stage-structured population model formulation

The model follows a simplified Anopheles gambiae life cycle with adult female mosquitoes laying eggs that hatch into early instars that develop into late instars. Late instars develop into pupae, which later emerge as adult mosquitoes. Each of these stages is then linked through or impacted by rates or probabilities such as fecundity, larvae, pupae or adult survival and development (Figure 1). The simplified version of the population dynamics was chosen to minimise sensitivity to the less-known parameters and processes while maintaining enough complexity to capture natural fluctuations in mosquitoes’ populations. Only female mosquitoes were modelled. Figure 1 illustrates the model. Below, a mathematical description of the model is provided, starting with transitions between life stages and then how those lead to population sizes at each stage at the next time point.

**Figure 1:**
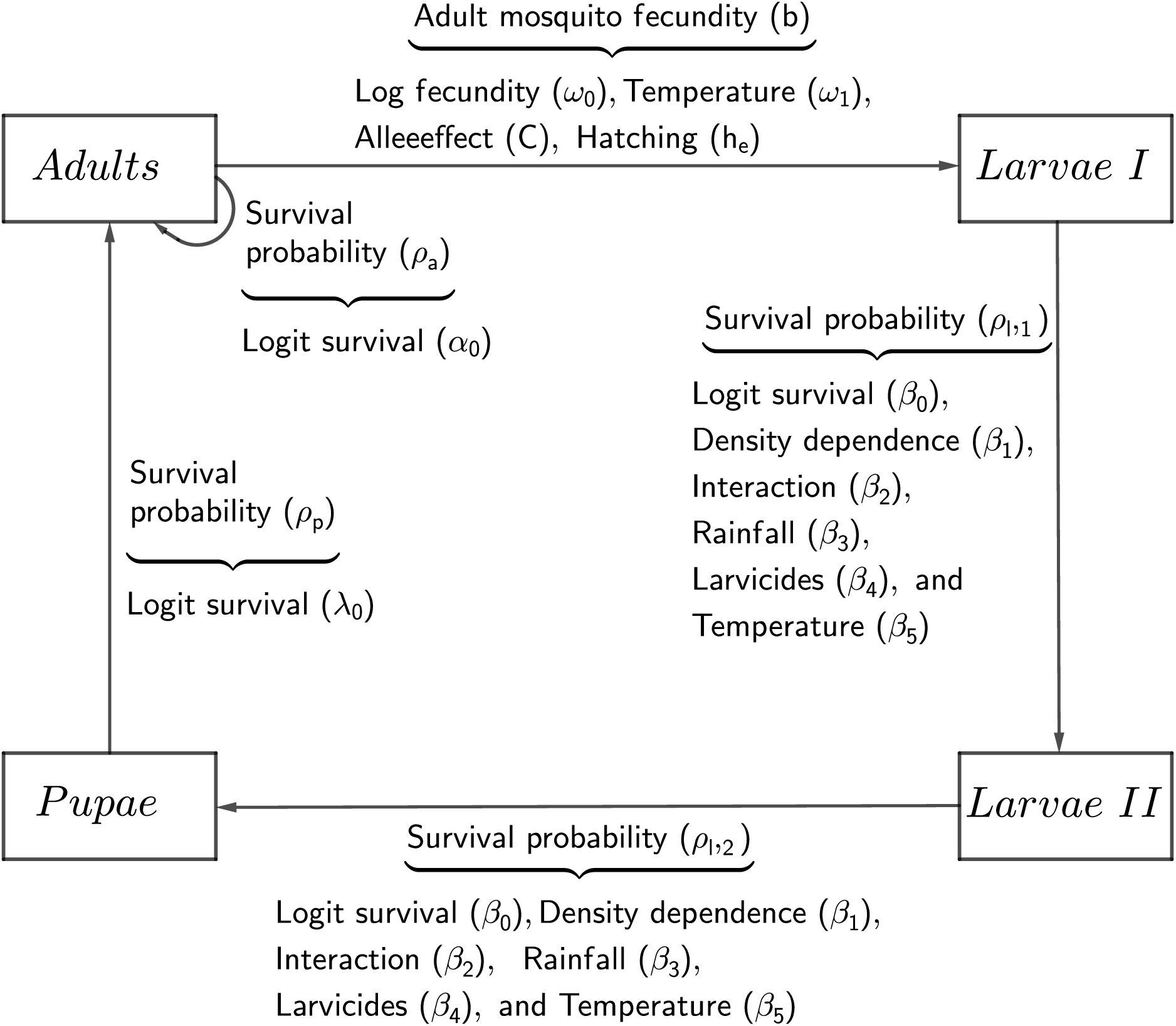
Schematic representation of the model showing stages of the mosquito life cycle together with the predictors of the life-history traits. Arrows correspond to the transition from one stage to another. Larvae I correspond to early instar larvae, while Larvae II correspond to late instar larvae.

### Transitions between life cycle stages

Here, eggs do not have their own stage, they are just a transitional state of early instar larvae. Briefly, the mosquito life cycle begins with eggs hatching into early instar larvae, N_l1_, to the late instar larvae, N_l2_, and then to the pupae, N_p_, who then become adults, N_a_ (Figure 1). To determine the total number of male and female mosquitoes in the population, a 50:50 sex ratio was assumed. To model the female mosquito population only, the total number of mosquito eggs laid per week (total fecundity) was halved, where only the proportion, h_e_, hatches to early instars.

The larval survival probabilities, 𝜌_x_ (where x refers to either early instar larvae, 𝑙_1_, or late instar larvae, 𝑙_1_) and number of individuals 𝑛 (i.e., early or late instar larvae), are defined by binomial distributions, with linearised functions 𝑆_x_, at ward 𝑖, and week 𝑡, through an inverse logit transformation such that,

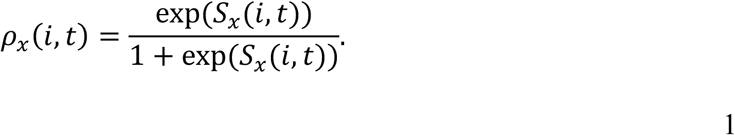

Survival probability of early instars, 𝜌*_l_*_1_, was defined through an inverse logit transformation of a linear function 𝑆*_l_*_1_, described as a function of the total larval abundance (𝑁*_l_*_1_+𝑁*_l_*_2_), rainfall (𝑅), larvicides (𝐿𝐴) and temperature (𝑇) such that:

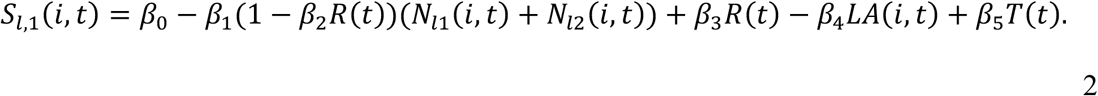

where 𝛽_0_is the mean logit (i.e., log odds) of larval survival when all covariates are 0 (i.e., centred rainfall = 0 mm, larvicide = 0 and centred temperature = 0 ^0^C). Temperature and rainfall were centred at their mean values of 27 ^0^C and 56 mm, respectively. That is, their means were subtracted so that each centred variable has a mean of zero, such that the intercepts of the survival and fecundity functions can be interpreted as logit survival probability conditional on mean temperature and rainfall. Parameters 𝛽_1-2_ describe the impact of DD on early instar larval survival, which is mainly regulated by rainfall and larval abundance. Specifically, 𝛽_1_is the effect of total larval abundance (i.e., 𝑁*_l_*_1_ and 𝑁*_l_*_2_) on early instar larval survival and here assumed to regulate the magnitude of DD. 𝛽_2_is the term that allows negative density-dependence to reduce when rainfall increases, which results in less larval competition. 𝛽_3_ is the effect of centred rainfall (𝑅) on early instar larval survival. Parameters 𝛽_4_, and 𝛽_5_ are the effects of larvicides (𝐿𝐴) and centred temperature (𝑇) on early instar larval survival, respectively.

The survival probability of late instars, 𝜌*_l_*_2_, was defined through an inverse logit transformation of the linear function 𝑆*_l_*_2_, and was structured similarly to Equation **1**. The late instar linear function was similar to the linear function of early larval instars (Equation **2**) (i.e., 𝑆*_l_*_1_ = 𝑆*_l_*_2_), hence, the parameter descriptions and values in functions 𝑆*_l_*_1_and 𝑆*_l_*_2_were similarly used in this study. Since the larval stage lasts for two weeks, it was divided into two stages, each lasting for one week (i.e. the time step of the model). Throughout the text, DD (i.e., 𝛽_1_) is regarded as a common parameter among the two larval stages.

Survival probability of pupae 𝜌_p_in ward 𝑖 at week 𝑡 was defined (as in Equation **1**) through an inverse logit transformation of the mean logit pupal survival (𝜆_0_). Adult survival probability 𝜌_a_in a ward 𝑖 at week 𝑡 was defined through an inverse logit transformation of the mean logit adult survival (𝛼_0_).

Mosquito fecundity was modelled as a Poisson process with mean 𝑏 in ward 𝑖 and week 𝑡 and linearised through an exponential transformation with temperature (𝑇). Fecundity is then regulated by the Allee effect parameter 𝐶, such that:

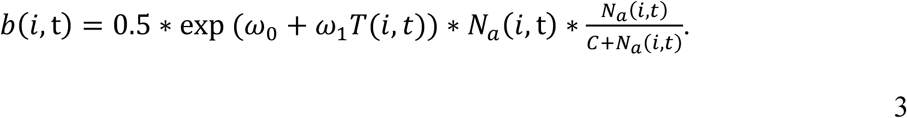

where 𝜔_0_is the mean log per capita fecundity when the centred temperature is 0 ^0^C, and 𝜔_1_ is the weekly effect of temperature on fecundity. AE is set as the probability of mating, where the number of female adults, 𝑁_a_, is scaled with a constant 𝐶. Specifically, the greater the value of 𝐶 relative to 𝑁_a_, the lesser the mating probability, leading to high impacts of AE (e.g., the mating probability will be less than 50% when 𝑁_a_ < 𝐶 and close to 100% when 𝑁_a_ ≫ 𝐶).

### Population size updates

The resultant survival probabilities (Equation **1**), functions (Equation **2**) and the total fecundity (Equation **3**) were used to obtain population abundances (which were used as one of the main outcomes) for both larvae (Equations **4** & **5**), pupae (Equation **6**) and adult mosquitoes (Equation **7**). Note that the survival probability will also incorporate the development rate, meaning that all surviving individuals move to the next stage.

Specifically, abundances of early instar larvae (Larvae I) were from a Poisson distribution with mean ℎ_e_ ∗ 𝑏 such that:

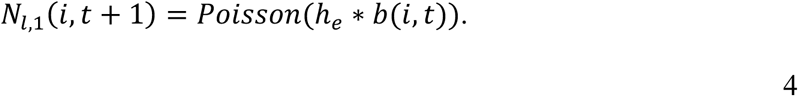

Abundances of late instar larvae (Larvae II) were from a binomial distribution with 𝑁*_l_*_,1_ trials and probability 𝜌*_l_*_,1_ such that:

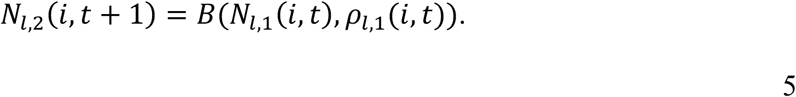

Abundances of pupae were from a binomial distribution with 𝑁*_l_*_,2_ trials and probability 𝜌*_l_*_,2_ such that:

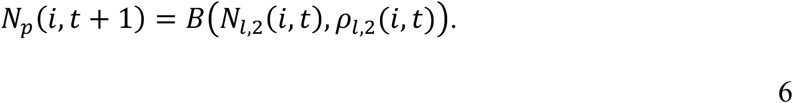

Total abundances of female adult mosquitoes were from binomial distributions with 𝑁_p_ and 𝑁_a_ trials and probabilities 𝜌_p_ and 𝜌_a_, respectively, such that:

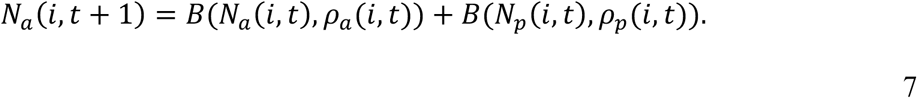

Probability 𝜌_a_ defines adults that remain in the system after surviving, and these individuals join those transitioning into adulthood from the pupal stage.

### Parameterization

The parameters in Equations **2** and **3** were given mean values based on literature, as shown in Table 1. The exceptions are the values for the parameters describing the AE (𝐶) and the interaction between DD and rainfall (𝛽_2_) because these are not available from the literature. Here, 𝐶 = 360 (which is number of female adults per thousand larvae and resultant percentage reduction in mating probability) was chosen as an intermediate value because it provides a baseline from which to explore increases and decreases in values of both AE and DD parameters. While 𝛽_2_ = 0.018 generates low population size (required for AE to occur) with stable population dynamics in the absence of an intervention, i.e., an average of 700 female adult mosquitoes (since the relevant size of local mosquito population is unknown, 700 was used as assumption but it could be as large as 10,000 or 100,0000, see Supplementary Information for results on ∼10,000). The value of 𝛽_2_ is fixed throughout the simulation, so DD will be controlled with 𝛽_1_. Since one important feature of the AE is its potential increase in stochastic extinction, the model was iterated 100 times (1000 iterations generated similar results, so 100 iterations were kept for efficiency), and the probability of extinction and the quantiles of the weekly adult mosquito abundances were estimated.

**Table 1:** Parameter definitions and values used in the simulation model. Only weekly estimates were used during the simulation. Original/literature values were converted to weekly values (methodology in SI). Fecundity (exponentiated from log scale) and survivals (inverse-logit transformed) are presented in biological units, rather than in their log or logit forms.

| Parameter | Parameter descriptions | Literature estimates<br>(Biological values) | Estimates<br>per week | Reference |
| --- | --- | --- | --- | --- |
| $\beta_0$ | Logit larval survival at mean rainfall and temperature in the absence of larvicide | 0.79-0.98 per day | 0.46 | [24,28,29] |
| $\beta_1$ | Effect of DD on larval survival, i.e., DD per thousand larvae | 5.5e-05 per week | 5.5e-05 | [24] |
| $\beta_2$ | Effect that allows DD to be weakened when there are abundant rainfall and therefore less larval competition | N/A | 0.018 | - |
| $\beta_3$ | Effect of rainfall in mm per day on logit larval survival | 0.0025 per 11 days | 0.0015 | [30] |
| $\beta_4$ | Effect of larvicides on larval survival | 0.057 per day | 0.25 | [31] |
| $\beta_5$ | Effects of temperature on larval survival | 0.0043 per 3 days | 0.01 | [32] |
| $\omega_0$ | Log fecundity at mean temperature | 50-300 per cycle | 182 | [33,34] |
| $\omega_1$ | Impact of temperature on the fecundity | - | 0.01 | - |
| $\lambda_0$ | Logit pupal survival at mean temperature | 0.7-0.8 per 2 days | 0.4 | [28,29] |
| $\alpha_0$ | Logit adult mosquito survival when temperature is 0 °C | 0.83-0.95 per day | 0.5 | [35,36] |
| $h_e$ | Eggs hatching rate | 0.23 per week | 0.23 | [37] |
| $C$ | Strength of mate finding AE, i.e., number of female adults per thousand larvae | - | 360 | - |

### Role of DD and AE in regulating mosquito populations

To address specific objectives, DD (𝛽_1_) and AE (C) were varied as follows: First, to illustrate the potential impacts on the population dynamics trajectories of these regulatory mechanisms in isolation, DD and AE were compared with low, medium and high levels. The 3 different values (low, medium and high) set for DD and AE are shown in Table 1, with medium levels (DD2 and AE2) corresponding to values that provide stable population dynamics. When exploring the impacts of DD, AE was kept constant either at 0 (absence of AE) or at its medium level (AE2). For exploration of AE, DD was kept at medium levels (DD2). The values for DD were defined from DD2, i.e., the literature value of 5.5e-05 (Table 1; DD2; [24]), where this value was halved to obtain a lower DD1 (𝛽_1_= 2.75e-05) and increased fourfold to a higher DD3 (𝛽_1_=2.2e-04). These values were chosen to illustrate the types of extreme impacts that high and low levels of DD might have on the population dynamics. Similarly, the AE was initially set at C=360 as mid-level (AE2, Table 1) and then halved to low AE1 (C=180) and increased twofold to high AE3 (C=720). AE was increased twofold as opposed to the fourfold increase in DD because when present, the impacts of AE on population dynamics are stronger, leading to extinction by doubling the initial C. Second, to explore their joint effects on population growth rate and probability of extinction, the three levels of DD and AE were varied simultaneously. Third, given the unknown values for these parameters, the percentage of population change was explored by investigating DD and AE in isolation across a range across the range of values from 90% reduction (>zero effect) of DD2 and AE2 to 100% increase (double effect for AE) or 300% (quadruple effect for DD) increase. This upper value is 2.2e-04 relative to AE3 and DD3. The percentage of population change was computed as the average across 100 independent simulations. To isolate baseline population dynamics and avoid confounding with treatment effects, all larvicidal coefficients in Equation **2** were set to zero, indicating the absence of intervention.

Growth rate and percentage of population change are distinguished in a way that the former is multiplicative, reflecting the compounding nature of mosquito reproduction in successive generations, while the latter is additive, providing a straightforward comparison between two population sizes at discrete time points. Growth rate is used when modelling population dynamics over time, as it captures the exponential potential of mosquito populations, while percentage change is used when evaluating the impact of a process or intervention, as it offers an intuitive measure of reduction or increase relative to a baseline [25,26].

### Impacts of DD and AE on sustained and short-termed interventions

The influence of DD and AE on population change when an intervention is applied either in a sustained (i.e., once application starts at week 48, it is continuously applied until the end of the simulation at week 193) or short-termed (i.e., an extended single short application lasting 7 months or two short applications lasting 3 months each, separated by 3 months) way was investigated. A short-term intervention regime was chosen because it is common for larvicide to be implemented for only a few months (e.g., 4 or 5). In contrast, the sustained intervention regime is not commonly used due to the high cost of resources, but serves to illustrate whether the intervention would have the potential to eliminate the population by preventing the population from bouncing back after suppression. For this, the parameters 𝛽_1_ and 𝐶 were set as described in the previous subsection, but here the larvicidal coefficient in Equation **2** was set to a non-zero value (as shown in Table 1). To illustrate the impacts of the intervention on the population dynamics, DD2 and AE2 were varied to explore the wider trade-offs between intervention and DD or AE by exploring the probability of population extinction through varying DD or AE and intervention from 90% reduction to 100% increase. The probability of extinction was computed as an average across 100 simulations.

### Trade-offs between negative density dependence and the Allee effect

Since the real values of DD and the AE are unknown, their effect sizes were varied simultaneously from 90% reduction to 100% increase on medium values (DD2 and AE2) set in Table 1. Population growth rates were estimated as the final divided by the initial mosquito population sizes. The growth rates were then averaged over 100 simulations, i.e., the simulation was iterated 100 times, recording the average growth rate over these simulations for each combination of DD and AE. A heatmap was used to show the population growth rate across wards for each percentage change in DD and AE. For illustration, the simulation was repeated under four intervention regimes: (a) without an intervention, (b) double short-termed, (c) single short-termed and (d) sustained intervention. A similar process as described in this subsection was repeated to compute the probability of extinction (SI).

### Initialization and implementation

The initial population size for both larval stages was set at 1000 larvae, while for the pupal stage, it was 900 pupae, and for the adult stage, it was 800 mosquitoes. These initial values were chosen based on the effect of 𝛽_1_, with a value of 5.5e-05 (Table 1) estimated from an abundance of 1000 larvae (see SI for results on large population sizes, e.g., 10,000 larvae).

The model was implemented in R software version 4.3.3 [27], and the R code is available in SI.

## Results

The normalized total mosquito population dynamics generated in the simulation model matched the dynamics of the observed mosquito population from the larvicidal intervention in Dar (Figure 2). The seasonal patterns characterized by peaks during rainy seasons and troughs in the dry seasons, including timing of the seasons, were generally captured by the simulation model; which is evident when comparing the simulated (Figure 2, red) with observed (Figure 2, black) abundances over weeks across wards. However, the model underpredicted the magnitude of peaks around weeks 50-55 and 155-160, whereas it overpredicted abundances during weeks 84-91. These under- and overprediction discrepancies are evident in Figure 2b, where several highest normalized numbers (>0.5) of observed mosquitoes fall below the 1:1 line, whereas very few normalized highest numbers (>0.6) of simulated mosquitoes lie above the line.

**Figure 2:**
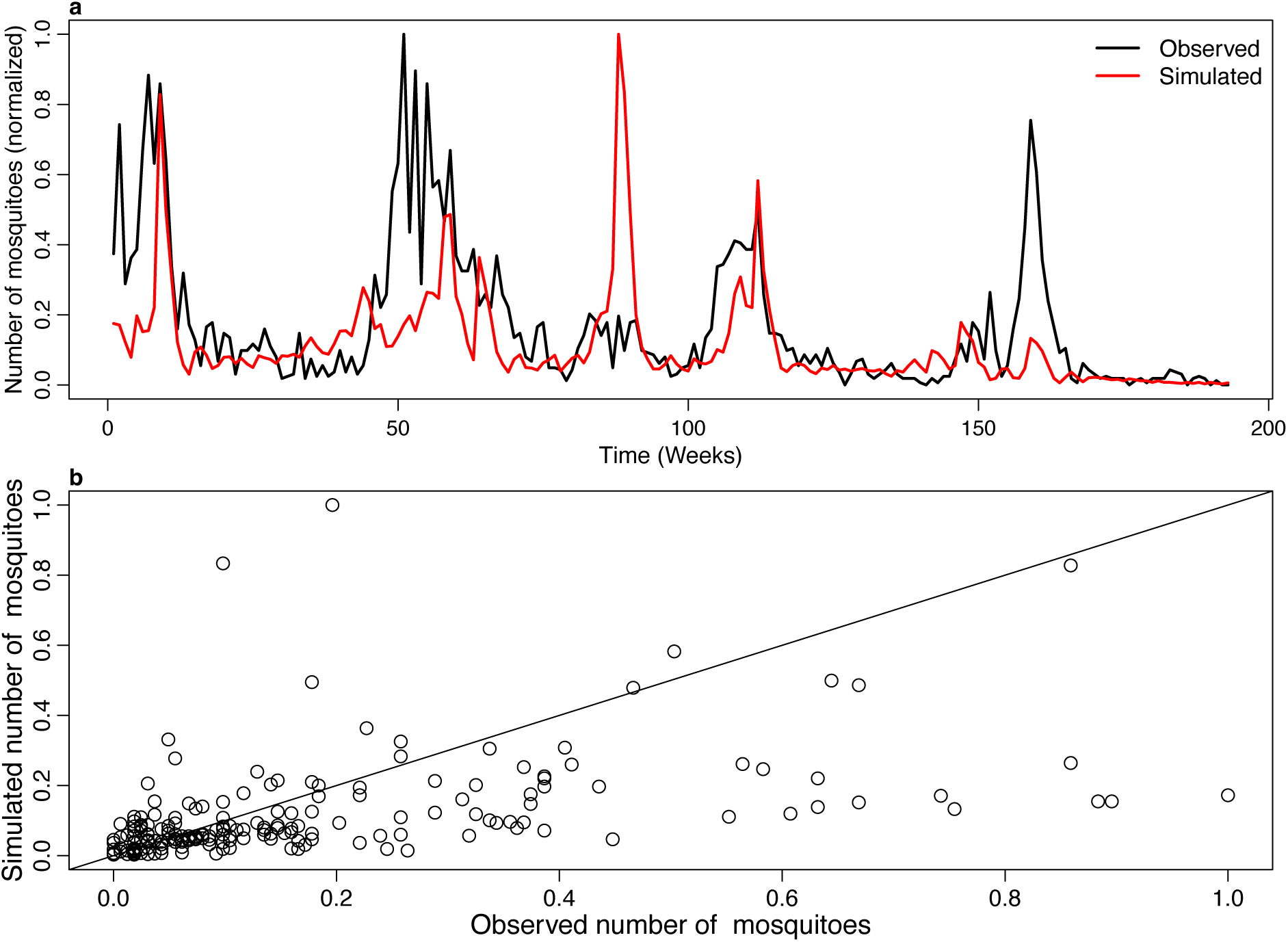
Model validation. (a) Reconstruction of the total mosquito population abundances (normalised) across wards, with observed mosquitoes (black) compared against the simulated trajectory (red) over weeks. (b) Comparison of observed versus predicted normalised total mosquito abundances across wards, where the 1:1 line diagonal indicates the goodness of fit.

### Role of DD and AE in Regulating Mosquito Populations

In isolation, neither regulatory mechanism at low or high levels impacted the population size in the long term (Figure 3). However, when varied together, especially at high levels, population decline accelerated. On the other hand, the population size stabilised at higher values as DD decreased. Specifically, DD3 with the AE2 resulted in a 100% reduction (i.e., extinction) in the mosquito population after the 37^th^ week (Figure 3a, black and blue lines). This reduction seems to be driven by AE since at DD3 without AE (i.e., AE=0), the population declined 90% after initialisation but soon stabilised again to an average of 180 mosquitoes and did not decline to extinction by the end of the simulation (Figure 3a, red line).

**Figure 3:**
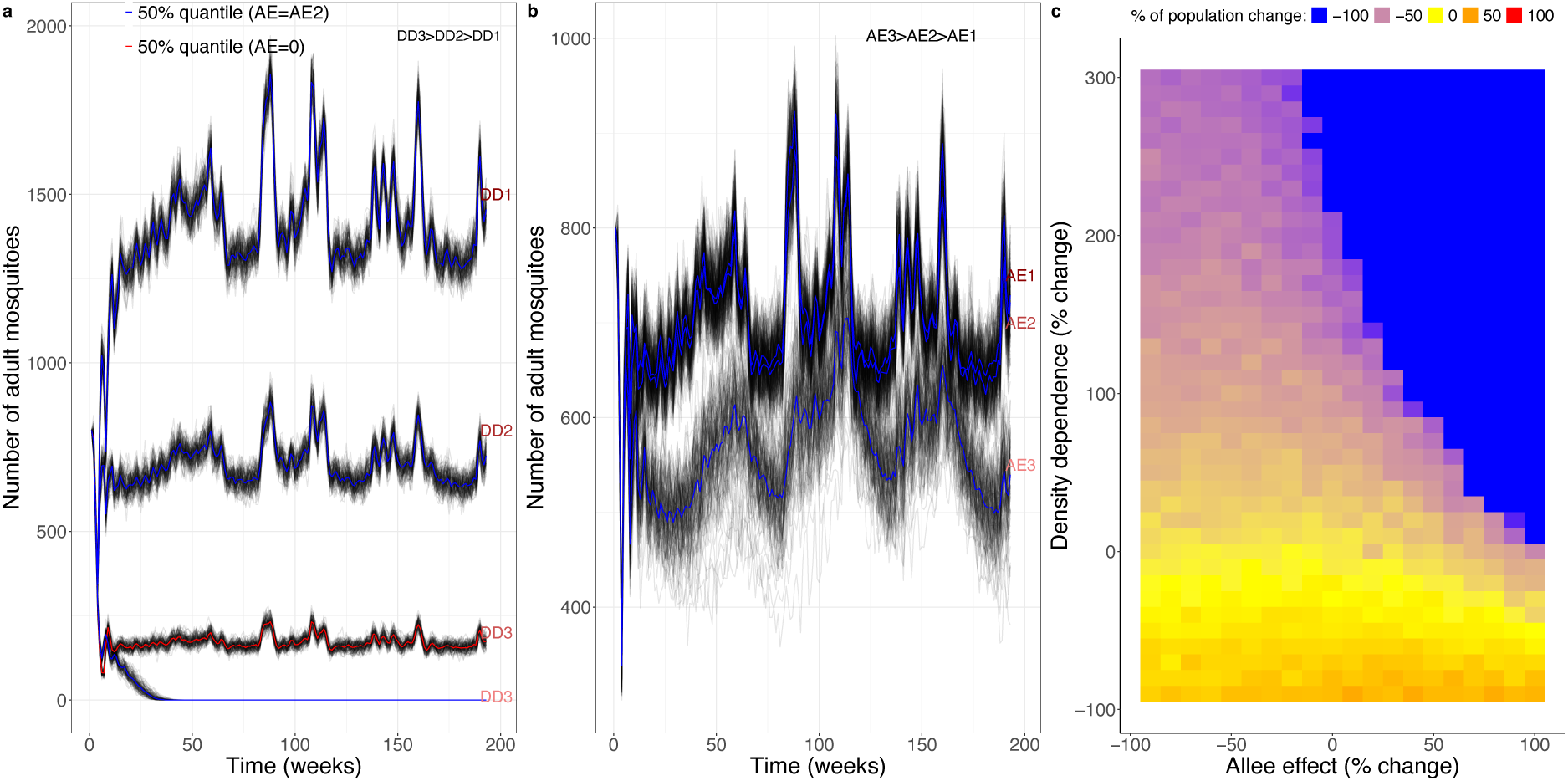
Role of DD and AE in regulating mosquito populations (without a larvicidal intervention). (a) Three DD levels (DD1=2.75e-05, DD2=5.5e-05 and DD3=2.2e-04) were used, keeping AE constant at 360 (AE2), and (b) three AE levels (AE1=180, AE2=360 and AE3=720) were used, keeping DD constant at DD2=5.5e-05. The red line in (a) corresponds to the 50% quantile of simulations with DD3 when AE=0, and the blue colour in (a) and (b) to the 50% quantile when AE=AE2. (c) Percentage of population change (relative to the initial size of 800 mosquitoes) after 193 weeks, with AE2 and DD2 varied across a range of values from 90% reduction to 100% or 300% increase. The percentage of population change was averaged over 100 simulations.

When varying AE, results were relatively similar in terms of average population size (Figure 3b). With AE1 and AE2 (with fixed DD2), population dynamics initially decreased 7% and 6.9%, respectively, before populations stabilised at an average of 700 mosquitoes (Figure 3b, black or blue lines). At AE3 (with fixed DD2), the population initially declined by 41.7% before stabilising at an average of 569 mosquitoes (Figure 3b, black or blue lines). Without DD (i.e., DD=0), population size increased and exploded irrespective of AE levels and hence not shown in Figure 3 in SI, indicating that DD is the regulatory mechanism that stabilises population size.

Broadly, when DD doubled and AE increased by 50%, population size quickly declined to extinction (Figure 3c, blue colour). Trajectories for DD1&2 at AE=0 are not shown here because they have the same dynamics as DD1&2 at AE=360, and for the same reasons, other combinations (i.e., DD1&2 vs. AE1&2) are not shown.

### Impacts of DD and AE on sustained and short-term interventions

As the duration of intervention increased, the mosquito population decreased in all studied scenarios, but populations only tended to go extinct in the presence of the AEs. With constant AE2 and short-term interventions (i.e., one or two short applications), the mosquito populations suffering DD1 and DD2 were initially reduced by 39% to 75% following each application, but bounced back to similar or higher levels. However, with DD3, the population declined to extinction (Figure 4c) after the first application (Figure 4e). Similarly, populations with AE1 and AE2 (with DD2) undergoing one or two short interventions declined by 39% to 99% before bouncing back to similar or higher levels (Figure 4d-f). In contrast, at AE3, population size declined to extinction, and there was no rebound regardless of the number of interventions.

**Figure 4:**
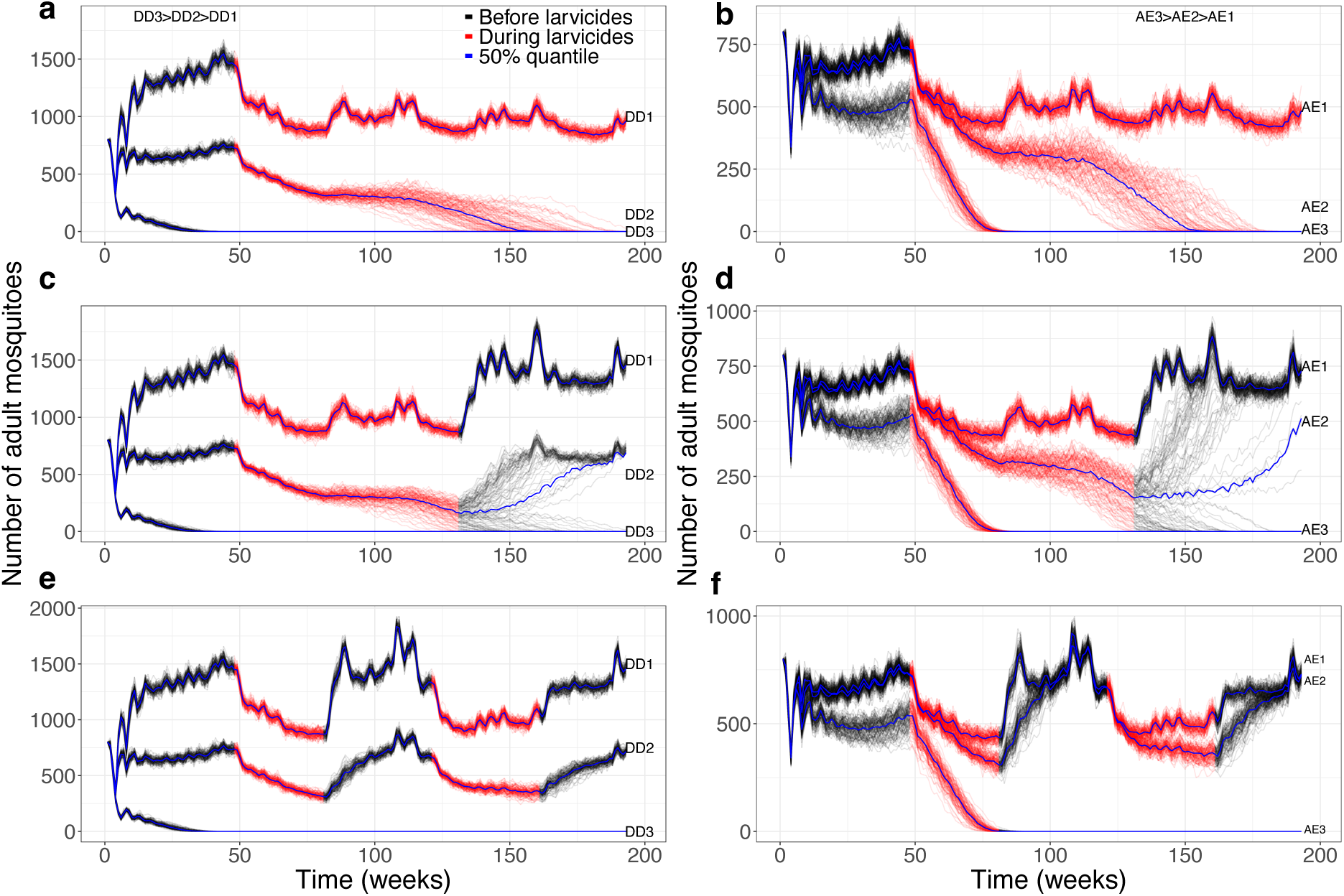
Impact of intervention on mosquito dynamics. Left column shows Mosquito populations regulated by DD at levels a) 2.75e-05, c) 5.5e-05 and e) 2.2e-04, while setting the AE constant at AE2=360. The right column shows the mosquitoes population with AE at levels b) 180, d) 360 and f) 720, keeping DD constant at DD2=5.5e-05. Larvicidal treatment was applied in three regimes: sustained larvicidal application (top row), a single short application (middle row) and two short applications (bottom row) and is represented in red.

When larvicide was applied in a sustained way from week 48 to the end of the simulation in week 193, the mosquito populations behaved in a similar way, but the strength of DD or AE required to drive populations to extinction was lower. For example, with AE2 at DD1, populations were able to bounce back and stabilise at an average of 1042 mosquitoes throughout the intervention period; however, at DD2 and DD3, the population declined to extinction (Figure 4a). At constant DD2 and AE1, the mosquito population declined but stabilised by the end of the simulation, while at AE2 and AE3, all populations went extinct (Figure 4b).

Overall, the probability of extinction increased with an increasing larvicidal effect, DD and AE (Figure 5a). Keeping the AE constant and varying DD and larvicidal effect across a range of values from 90% reduction to 100% increase, it was found that there was a need to increase DD and larvicidal effect by 50% to drive the population to extinction. While the patterns are similar for constant DD with varying AE and larvicidal effect, the probability of extinction never reached 0 (ranging between 0.25 to 1, Figure 5b) and increasing AE by more than 50% was sufficient to drive the population to extinction even with minimal larvicidal effect. Together, these results indicate that during intervention, the presence of even small AE levels would facilitate population control, despite the ability of the population to bounce back from DD.

**Figure 5:**
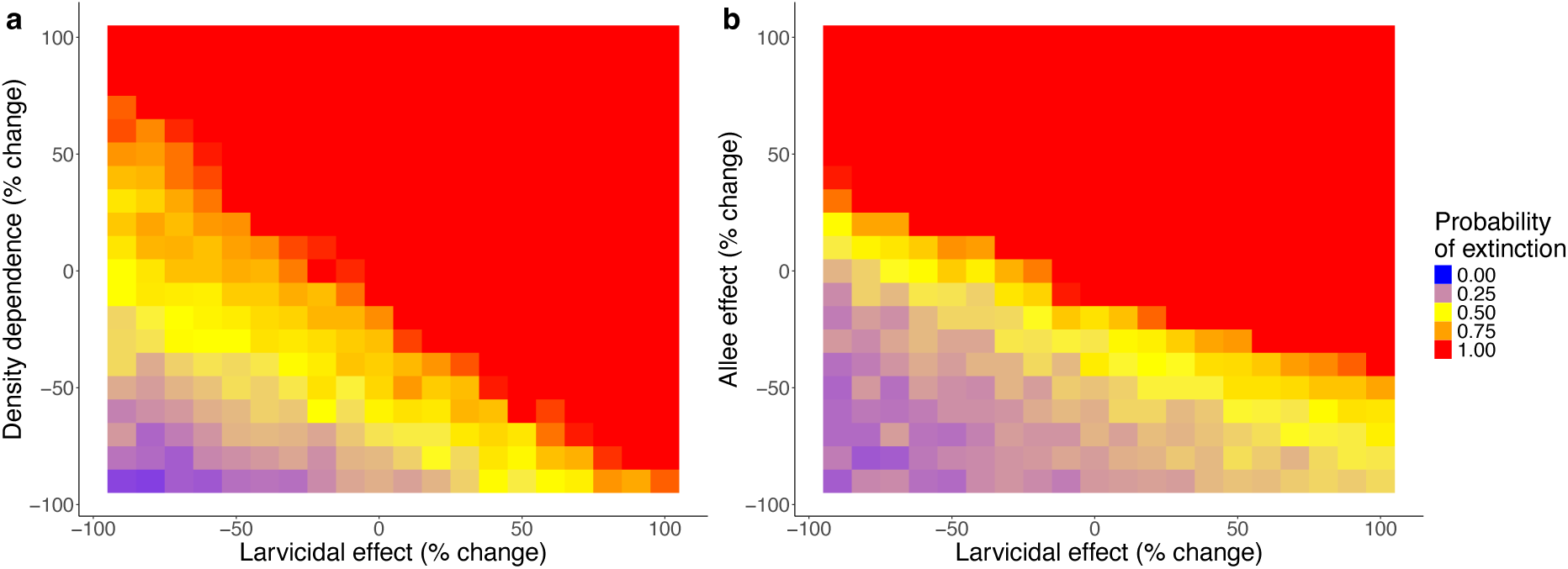
Heat maps showing the probability of extinction (averaged over 100 iterations) by week 193 after varying across a range of values from 90% reduction to 100% increase in sustained larvicidal effect size, 0.25, and (a) DD effect size, 5.5e-05, setting AE constant or (b) AE effect size, 360, setting DD constant.

### Trade-offs between negative density-dependence and Allee effect in the mosquito population regulation

As DD and AE increased, the population growth rates decreased, but AE tended to accelerate population decline (Figure 6), meaning that a small increase in AE leads to a bigger impact on the mosquito populations. For example, in Figure 6c & 5d, differences in patterns are not visible because AE had a bigger impact as it increased. While DD with less AE (i.e., 90% reduction) could not lead to population growth rates below one, higher levels of AE with less DD (i.e., 90% reduction) caused mosquito population rates to decrease below one, especially with a sustained intervention (Figure 6b-d). With lower levels of DD and AE (i.e., DD2 and AE2 reduced by 90%), the population size was high, resulting in growth rates above one, with or without larvicidal intervention (Figure 6a-d). Without intervention, the population growth rate could not decrease below one unless DD and AE were both increased by at least 50% (Figure 6a). With a double short application of intervention, only a 10% increase in DD and AE could drive the population growth rate below one (Figure 6b). Similarly, with a single short application of intervention, only keeping DD and AE constant could drive the population down at a growth rate below one (Figure 6c). In Figure 6d, the mosquito population growth rate declined to less than one without any change in DD and AE (i.e., 5.5e-05 and 360, respectively). However, with almost no DD, the population growth rate slowly declined below one if and only if the AE was doubled in the presence of single or double short applications of intervention (Figure 6b, c). When DD was increased by at least 50%, population growth rates declined below one even before reaching a 50% increase in AE if and only if there is an intervention (Figure 6b-d). When DD and AE were both increased by at least 50%, their combination drove the growth rate to zero (Figure 6b-d). However, the presence of a sustained intervention accelerated the combined effect of DD and AE to achieve zero growth rates (Figure 6d). Increasing AE substantially reduced the growth rate once the intervention was introduced; this explains why Figure 6c and 5d are nearly identical because the short-term intervention had already driven the population to extinction with only modest increase in AE.

**Figure 6:**
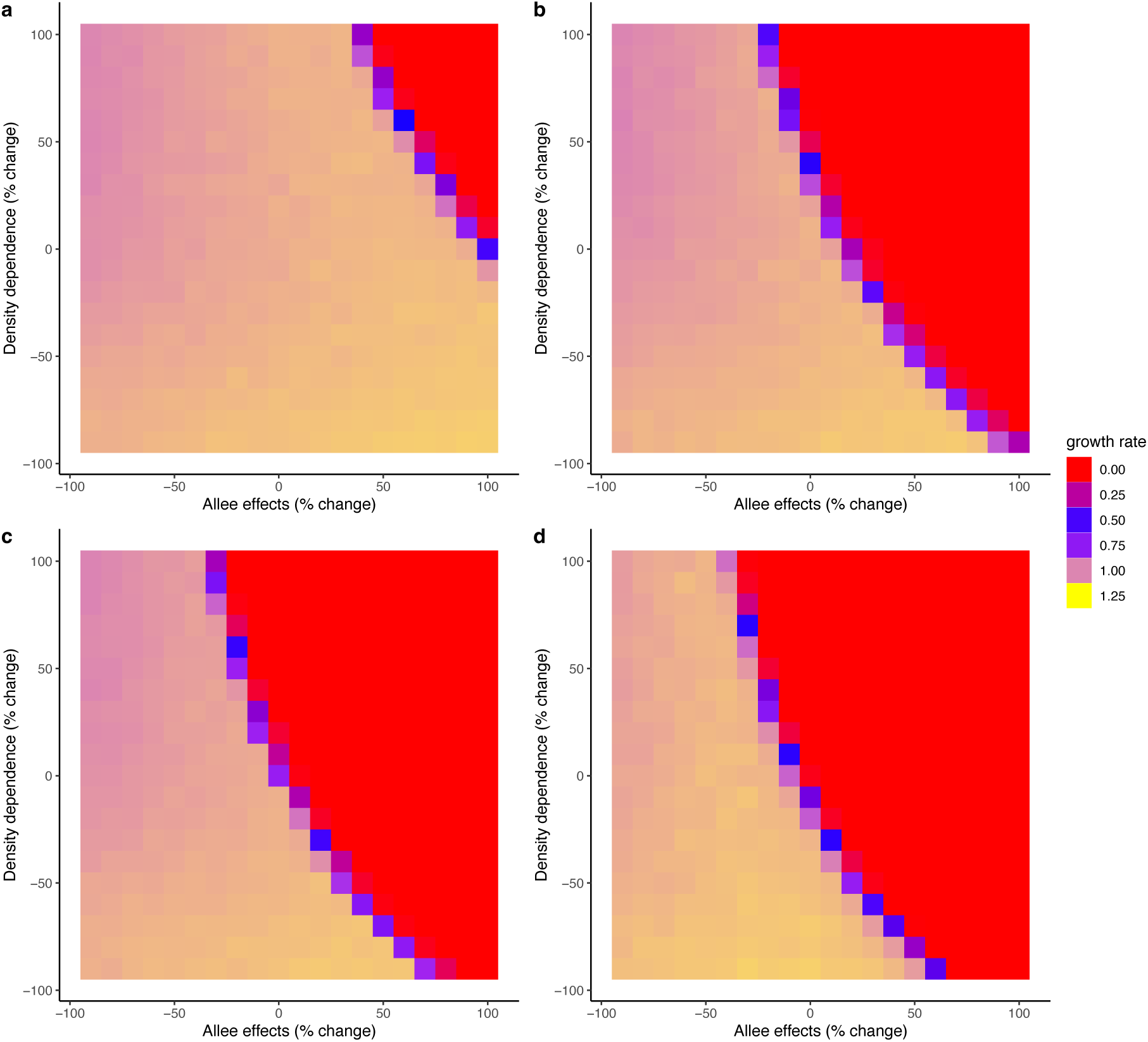
Heat maps showing the population growth rates averaged over 100 simulations for each percentage increase or decrease in DD and AE mean values, 5.5e-05 and 360, respectively; (a) without intervention, (b) with double short application of intervention, (c) single short applications of intervention and (d) with sustained intervention.

The model results and parameter values for DD and AE were relative to the population size of 1000 mosquitoes. This was to match the observed estimates from the case study in Dar. However, it is acknowledged that this size is small relative to a true field population and thus the sensitivity of the results to population size was also investigated. The results show that if DD and AE parameters are scaled for larger populations, the resultant patterns remain consistent. For example, when the population size is 10,000 mosquitoes, AE2=3600 and DD2=5.5e-06, the results (population trend) were broadly the same. See Supplementary Information for detailed results for a 10,000 total population size.

## Discussion

Understanding the interplay between vector control interventions and vector population dynamics is key to making informed decisions and accelerating disease elimination. In vector-borne diseases such as malaria, for which control is widespread, but populations tend to persist, dissecting the role of these regulatory processes (i.e., DD and AE) might help determine the type of intervention, effort (i.e., the intensity needed), duration and the timing over which the intervention should be deployed to prevent populations from rebounding. Neither DD nor AE alone was sufficient to substantially reduce the population, however, their combined effects accelerated the decline and ultimately drove the population to extinction. Sustained application of interventions could reduce the population sufficiently to ‘activate’ AE and increase or accelerate the likelihood of mosquito populations going extinct, but short-term application of interventions could lead to a population rebound driven by DD. This trade-off between DD and AE in the regulation of mosquito populations opens the possibility that vector control measures could require fewer resources to achieve large outcomes. To the best of current knowledge, this is the first study to systematically evaluate the importance of DD and AE and their implications for vector control broadly and specifically for malaria vector elimination, and it demonstrates that extinction can occur and be accelerated when the population size is small, and AE is strong relative to DD.

A mosquito population cannot grow indefinitely as the DD eventually regulates its growth [38,39]. Previous studies found that incorporating DD in models reflects more accurately the trends in the development and survival of immature mosquitoes, compared to those without DD [40–42]. Similarly, [13] reported that with higher larval density, survival declines (i.e., mortality increases) and the development time (time to reach pupal stage) increases, and in turn, there are fewer surviving larvae, experiencing less competition, making the population size increase again. This aligns with the results, where populations experiencing DD alone initially declined to lower- or mid-levels due to short applications of larvicides but subsequently rebounded to higher levels than before after the intervention had stopped. These findings suggest that interventions such as larvicides should be applied frequently and consistently to effectively reduce mosquito populations. Short-term interventions alone are unlikely to drive populations to extinction, as stopping treatment allows remaining mosquitoes to quickly recover due to reduced competition for resources.

While DD has been extensively studied [13,39,41,43–46], AEs and their implications for regulating small malaria mosquito populations remain largely overlooked to the point that their existence is not yet proven in the field. It is assumed that AE in mosquitoes can occur via mating limitation, here modelled by a reduction in the number of eggs produced. However, *Anopheles gambiae s.l.* mosquitoes exhibit a unique mating system known as lekking, where males establish fixed display territories, referred to as leks. In these areas, males perform displays to attract females, who then visit the leks and select mates based on the quality of the display [47]. While the assumption is still valid, this mating system may imply that one male can mate with many females, and hence AE could be either absent or difficult to achieve, potentially contributing to the rarity of mosquito population collapses. Regardless, this study demonstrates that if present, AE could have important implications for the effectiveness of common interventions such as larviciding. This is because interventions typically work by reducing population sizes to low levels, and it is shown that if AE kicks in, populations can decline much faster to extinction, even if the intervention level is set at a constant effort. Without AE, the population could recover quickly, making interventions non-sustainable. The only exception is when interventions are applied long-term, which unfortunately is often not feasible due to limited resources available to control VBD in low- and middle-income country settings.

Populations undergoing both DD and AE were the most susceptible to the impacts of interventions, with sustained interventions driving the population to extinction when subject to intermediate levels of DD and AE. Here, it means that DD was not sufficiently strong to overcome the AE when the population became small due to the intervention. It is important to note that the population undergoing DD or AE alone did not reduce the population to extinction, even when intervention was sustained for a long period. In other systems, *Bythotrephes longimanus* exhibited strong AE that impacted establishment success in low-density populations, particularly under demographic stochasticity and limited propagule pressure [48]. Strombus gigas also demonstrated AE with reproductive failure occurring below critical adult densities despite potential mates, highlighting mating success failure even under nominal adult abundance [49]. Given DD and AE, sustained intervention was more effective in driving the population to extinction than short-term intervention. Generally, given the AE, the probability of population extinction increased with the duration of the intervention. This implied that an intervention such as larviciding could take advantage of these main regulatory processes to accelerate malaria elimination and wrap up the endgame, though quantifying the threshold at which AE kick in remains a challenge and will likely be context dependent. Additionally, it is important to note that findings from this study imply that the scale of regulatory processes is crucial, whereby only high levels of DD and AE can lead to smaller population sizes and eventually population extinction.

The modelling framework developed here assessed the effect of DD and AE trade-offs on mosquito populations. The choice for DD and AE values is limited to 1000 adult mosquitoes, indicating that their values will need adjustment for population sizes exceeding 1000. The model produced population dynamics similar to those found in the field; however, there was underprediction restricted only to normalisation and not evident with raw (non-normalised) abundances. The normalisation outcomes were consistent with validation objective, which focused on evaluating whether the model captured the temporal dynamics of the observed data. Normalisation was employed to facilitate comparison across wards, and thus the discrepancies noted in the model reflect scaling rather than a failure to reproduce the underlying temporal dynamics. While most parameters were parameterised from semi-field or field studies of the same species and region (i.e. Anopheles gambiae from Tanzania), there were challenges due to a scarcity of experimental data [50]. This is particularly evident for the AE parameter, which has never been identified in the field. AE was modelled primarily as a reduction in fecundity, simplifying the broader ecological mechanisms in mosquito populations. Key factors such as reduced mating success or sex ratio imbalances at low densities were not explicitly addressed by the model. Assumptions regarding AE strength and population size carry uncertainties that influence the model’s predictions. Future work should explore alternative formulations of AE and empirically based parameters to better reflect population dynamics. Nonetheless, this framework can be adapted to identify AE in the field and fills a gap in the understanding of these regulatory processes and their implications for malaria vector control and elimination efforts.

## Conclusion

Understanding ecological dynamics that regulate mosquito populations is essential not only for vector control but also for advancing broader ecological theory. Anopheles mosquitoes, as selected species that exhibit a unique mating system, such as a lekking system, may suppress AE. Yet, exploring the potential for AE remains critical, especially when interventions push populations toward low-density thresholds. More importantly, AEs are increasingly recognised across taxa, from cooperative breeding in birds and social structure in mammals, to pollination and seed dispersal limitations in plants, and mate-finding limitations in insects and reptiles. Assessing these underexplored AE mechanisms can highlight both resilient and vulnerable stages in life cycles across ecological systems. Identifying where and when AE kicks in can inform control strategies that leverage these dynamics, whether to control disease vectors, manage endangered species, or control invasive populations.

## Declarations

### Ethics approval and consent to participate

Not applicable.

### Consent for publication

Not applicable.

### Availability of data and materials

Data on adult female *An. gambiae* mosquitoes used for model validation were obtained from previous studies (Geissbühler *et al.*, 2009; Maheu-Giroux and Castro, 2013). Environmental data used in this study are available in an open-source global weather data repository at https://globalweather.tamu.edu/#pubs (SWAT, 2022).

## Authors’ contributions

AMK, SSK, DTH, PCDJ and MV conceived the ideas and designed the methodology; AMK developed the model and performed the analysis with the guidance of SSK, DTH, PCDJ and MV; AMK, PCDJ and MV interpreted the model results; AMK drafted the first version of the manuscript. AMK, PCDJ and MV led the writing of the manuscript. All authors contributed critically to the drafts and gave final approval for publication.

## Acknowledgements

We are grateful to the members of the Data Science and Mathematical Modelling team, Entomological Technicians and Research Scientists at Ifakara Health Institute who provided their support to AMK during his visits to the institute. We are also grateful to Late Prosper Chaki, who provided a comprehensive explanation to AMK regarding the data from the large-scale larvicidal control in Dar es Salaam, Tanzania.

## Supplementary information

The supplementary information is included in a separate file named *Supporting information*.

## List of abbreviations

AE: Allee effects
Dar: Dar es Salaam
DD: Negative density-dependence
Larvae I: Early instar larvae
Larvae II: Late instar larvae
LA: Larvicides
SI: Supporting information

