## Supporting information for "Implications of the trade-offs between negative density-dependence and Allee effects for vector control"

**List of affiliations**

[1]School of Biodiversity, One Health and Veterinary Medicine, University of Glasgow, Graham Kerr Building, Glasgow G12 8QQ, United Kingdom

[2]Environmental Health & Ecological Sciences Department, Ifakara Health Institute, P.O. Box 78 373, Dar es Salaam, Tanzania

[3]The Pan-African Mosquito Control Association, KEMRI Headquarters, Mbagathi Road, Nairobi, Nairobi 54840-00200, Kenya

[*]Corresponding authors

 &

[§]Joint senior authorship

**List of email addresses:**

This Supporting Information file contains a simple review of the evidence for Allee effects in female adult *An. gambiae* mosquito data, temperature and rainfall data from Dar es Salaam, a list of tables for the combination of varied values between DD, AE and LA, plots and R code used to produce plots for the results section.

**A simple review of the evidence of the Allee effects in natural systems**

The occurrence and relative importance of DD and AE for population dynamics and stability have been demonstrated across taxonomic groups, including fish, reptiles, birds, mammals, insects, and plants [1,2]. For example, DD has been described in butterflies (*Maculinea alcon* and *M. teleius*), where DD alone explained up to 62% and 42% of interannual population variation, respectively, far more than the explanatory power of environmental drivers such as weather and land use (Nowicki *et al.*, 2009). For insects such as *Spodoptera exigua*, increasing larval density induced defensive responses in tomato plants, thereby reducing food quality and decreasing larval growth and survival [4]. AEs are less well described, but examples include new plant populations, where low numbers of plants can impede conspecific pollen transfer, potentially driving the population to extinction if founder plants are not enough to attract pollinators to regularly service plants growing close to them [5,6]. AEs have also been observed in vertebrates, e.g. a goldfish population reduces more rapidly at smaller sizes [7]. Similarly, experiments in mammals, including deer mice and red-backed voles, showed that high as opposed to low population sizes enhance reproduction, survival, and protection from threats [8,9]. In insects such as gypsy moths and beetles, AE mechanisms include mate limitations and predation [10–14], but examples are sparser. For example, [15] showed that *Callosobruchus chinensis* and *Tribolium confusum* can suffer mate limitations, which can lead to extinction at low densities. Importantly, these regulatory mechanisms have been used to support the management of species. In the United States, AEs facilitated the control of biological invasions through the establishment and spread of non-native species by limiting mating of invaders such as *Philornis downsi,* red turpentine beetle and *Dendroctunus valens* [16,17]. Given the potential opposite effects of DD and AE on efforts to control vector and pest populations, it is important to understand the trade-offs of regulatory mechanisms such as DD and AE, especially in small populations.

### ***Anopheles gambiae*** **field mosquito population sizes**

Figure 1 shows the monthly *An. gambiae* mosquito population abundances per ward and across all fifteen wards of Urban Dar es Salaam, Tanzania.

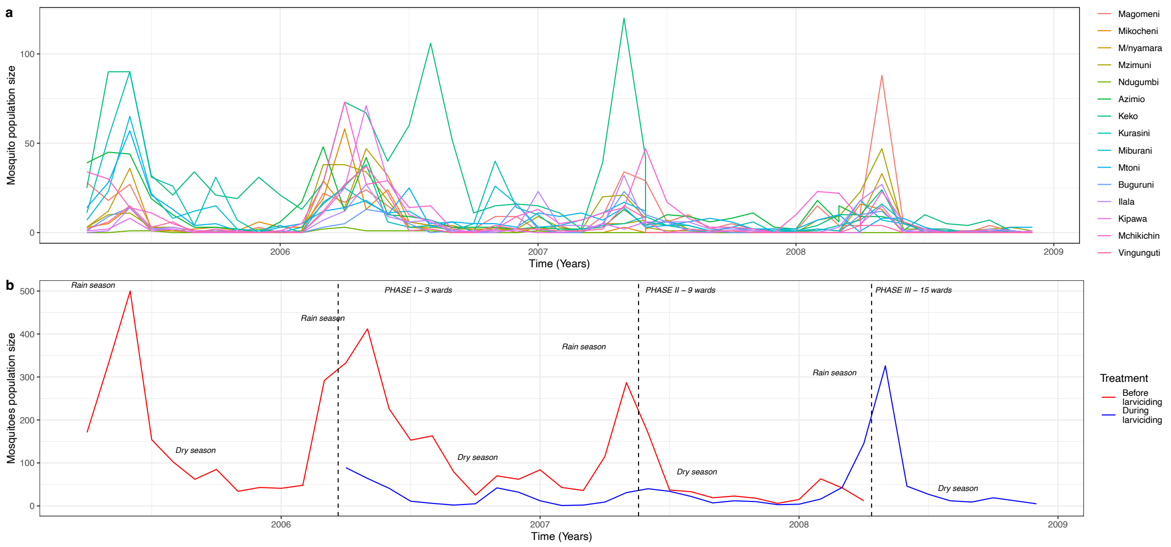


Figure 1: Adult female An. gambiae mosquito population abundances aggregated monthly from 2005-2009 (a) per ward and (b) across wards before and during large-scale larvicidal control in Dar es Salaam, Tanzania.

### **Weekly simulated mosquito population dynamics, non-centred and centred rainfall and temperature during the large-scale larvicidal control programme in Dar es Salaam**

**Figure 2** shows how the dynamics of data simulated with an intervention equal to zero match the dynamics of the centred and non-centred rainfall.

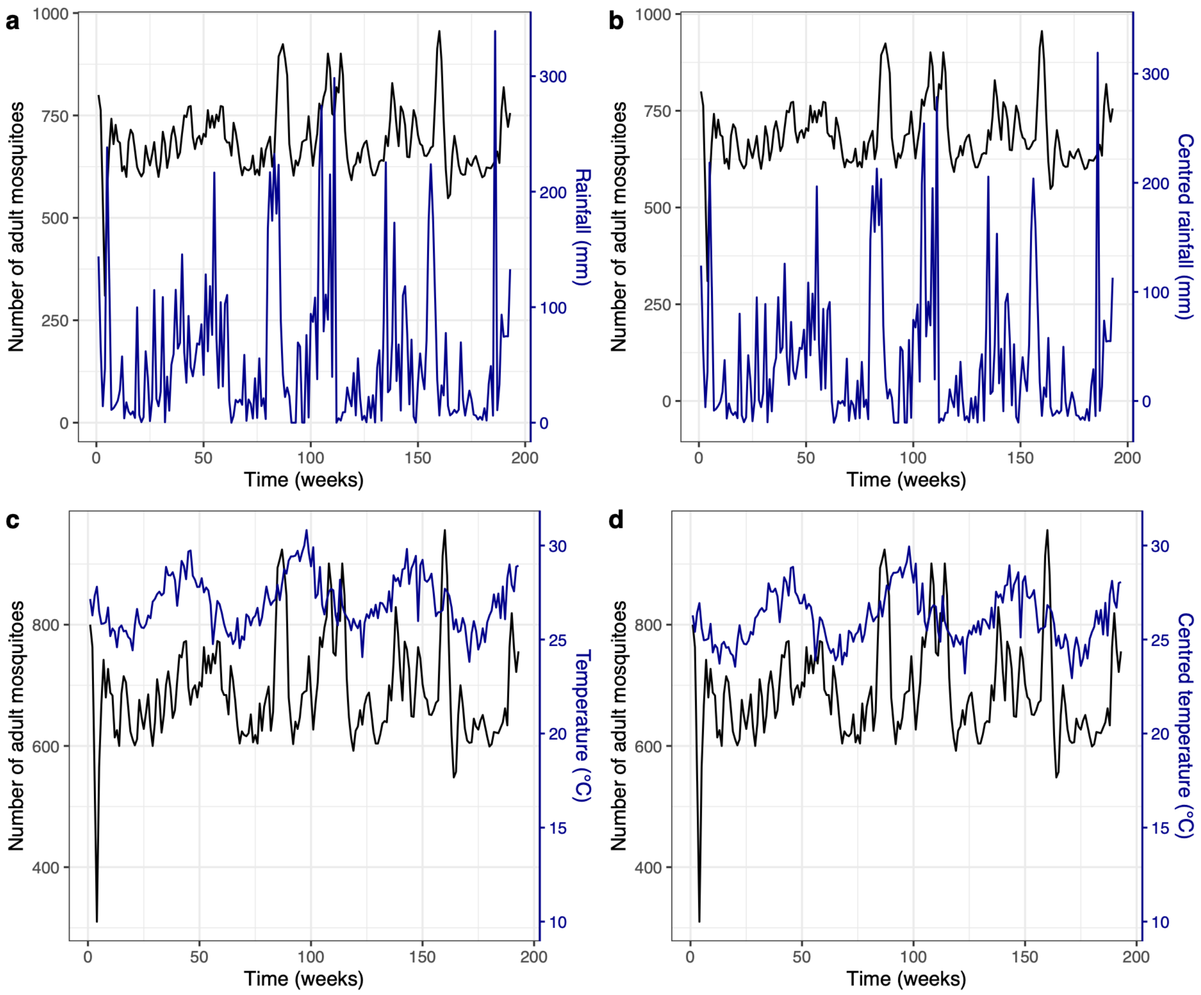


Figure 2: Weekly (a) non-centred rainfall and (b) centred rainfall (dark blue), together with (c) non-centred temperature and (d) centred temperature (dark blue), plotted alongside simulated population dynamics without intervention (black) during the large-scale larvicidal control in Dar es Salaam, Tanzania.

### Model validation

**Figure 3** shows a simulated trajectory from the simulation model capturing the observed trajectory from the large-scale larvicidal intervention in Dar es Salaam, Tanzania, as well as their 1:1 line indicating goodness of fit.

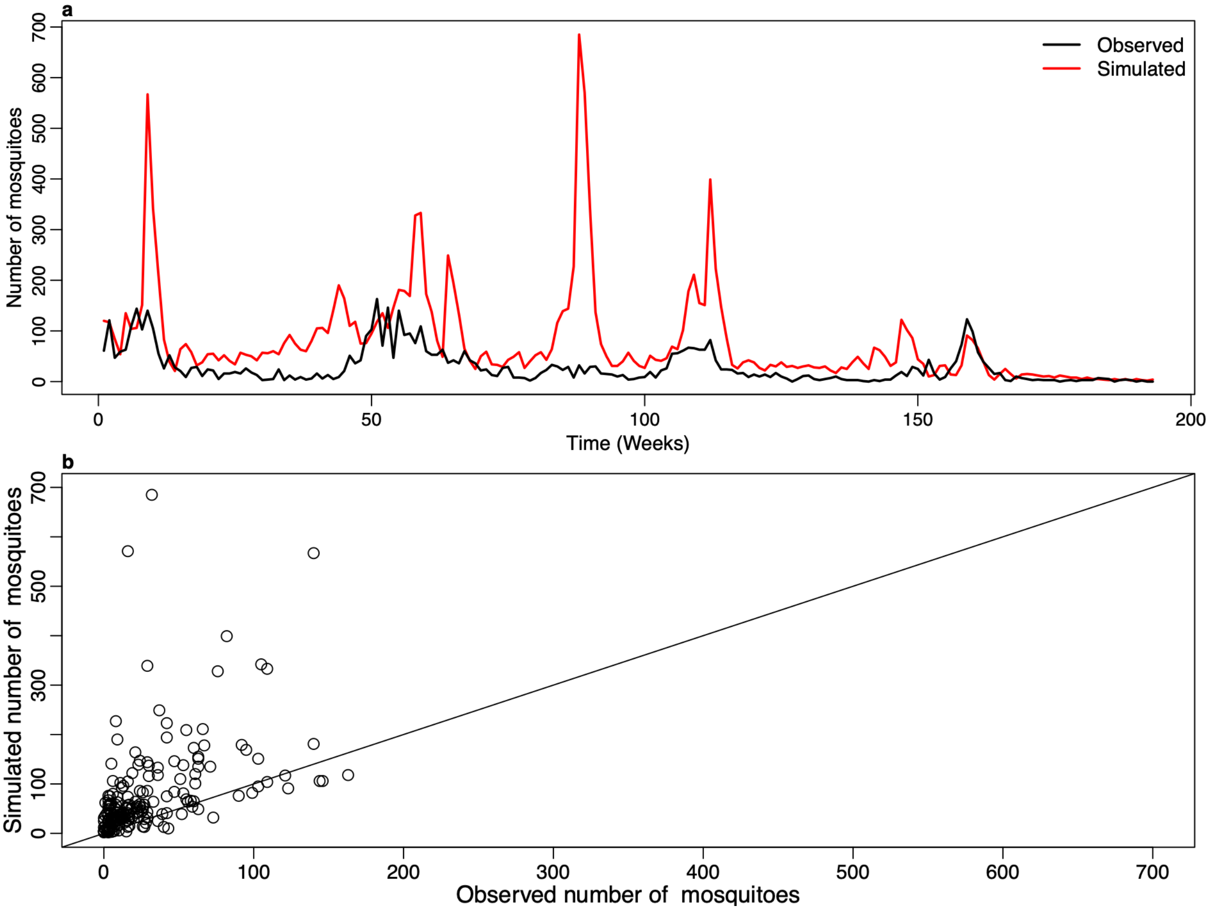


Figure 3: Model validation. (a) Reconstruction of the total mosquito population abundances (non-normalised) across wards, with observed mosquitoes (black) matched against the simulated trajectory (red) over weeks. (b) Comparison of observed versus predicted normalised total mosquito abundances across wards, where the 1:1 line diagonal indicates the goodness of fit.

**Table of selected combinations of density dependence (DD) and the Allee effect (AE) variations**

Table 1: Study variables and scenarios used for sustained and short-termed intervention

| **Study variables and scenarios** | **Simulated values** |
|---|---|
| Sustained larvicidal intervention | 48th to 193rd week |
| Short-termed larvicidal intervention (single short application) | 48th to 130th week |
| Short-termed larvicidal intervention (two short applications) | 48th to 80th and 121st to 150th week |
| Density-dependence levels i.e., DD1, DD2 and DD3 (when the Allee effect set to 360) | 2.75e-05, 5.5e-05, 2.2e-04 |
| The Allee effect levels i.e., AE1, AE2 and AE3 (when density dependence set to 5.5e-05) | 180, 360 and 720 |

All combinations of density dependence and the Allee effect variations (AE, DD)

| | | Negative density-dependence (DD) | | |
|---|---|---|---|---|
| | | 2.75e-05 | 5.5e-05 | 2.2e-04 |
| The Allee effect (AE) | 180 | (180, 2.75e-05) | (180, 5.5e-05) | (180, 2.2e-04) |
| | 360 | (360, 2.75e-05) | (360, 5.5e-05) | (360, 2.2e-04) |
| | 720 | (720, 2.75e-05) | (720, 5.5e-05) | (720, 2.2e-04) |

**Population without density dependence**

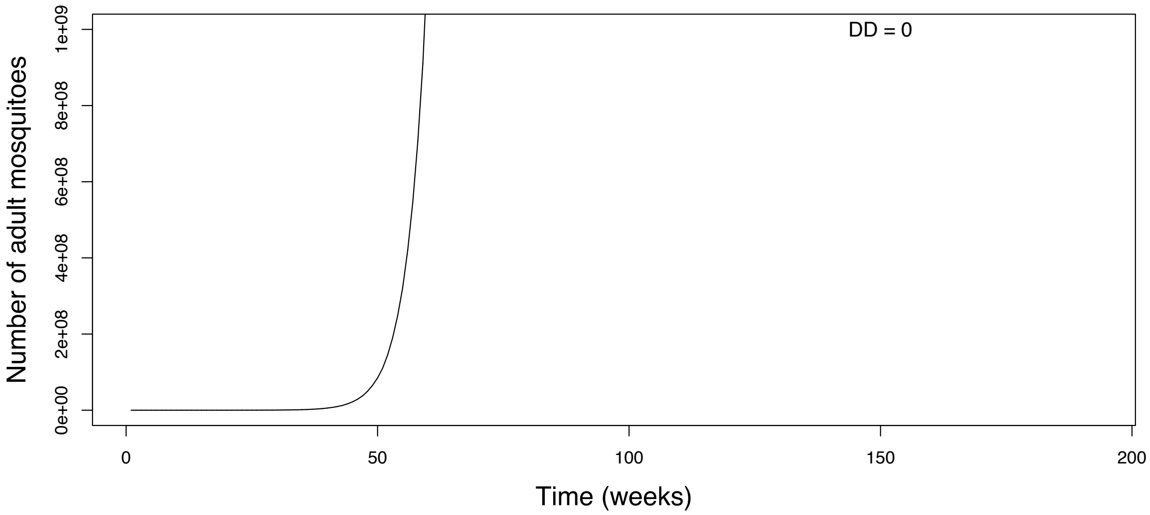


Figure 4: Population size explodes to infinity in the absence of density dependence (i.e., $\beta_1 = 0$).

**The Leslie Matrix for the estimation of adult mosquitoes' population growth rate**

The Leslie matrix method is commonly used in ecological studies to predict sizes and growth rates of stable stage-structured populations by using individuals' survival probabilities and fecundity rates in different life cycle stages [18]. In this study, the Leslie matrix was utilised to assess the growth rates of the stable adult mosquito populations based on the stage-structured population model developed in the methods section of the main text. Since it is difficult to observe the growth rate estimates directly from the simulation when the population stabilises, because most values will be centred around the same point, the Leslie matrix helps to visualise the deterministic growth rate as a smooth line. The results here help to understand how density dependence and Allee effects regulate the growth rate of a stable mosquito population over time.

When written in matrix form, $N(t+1) = LN(t)$ where $L$ is a Leslie matrix and *N(t)* is the population abundances at time t, the dynamics are given by:

$$\begin{pmatrix} N_{l,1}(t+1) \\ N_{l,2}(t+1) \\ N_p(t+1) \\ N_a(t+1) \end{pmatrix} = \begin{pmatrix} 0 & 0 & 0 & b_r \\ \rho_{l,1} & 0 & 0 & 0 \\ 0 & \rho_{l,2} & 0 & 0 \\ 0 & 0 & \rho_p & \rho_a \end{pmatrix} \begin{pmatrix} N_{l,1}(t) \\ N_{l,2}(t) \\ N_p(t) \\ N_a(t) \end{pmatrix}$$

**1**

The maximum eigenvalue of the matrix $L$ is the per capita growth rate of the population (assuming the population is at a stable stage structure), but in contrast to the simple density-dependent Leslie matrix, the parameters in Equation 1 are density dependent.

The per capita population growth rate was also obtained directly from the simulation model as an average across 100 simulations using the abundance of adult mosquitoes in the formula below.

$$Gr(t) = \frac{N(t+1)}{N(t)}$$

Whereby $Gr(t)$ is the population growth rate.

The per capita growth rates were initially computed using the Leslie matrix, and then compared with those generated directly at each time step of the simulation model. A 5% proportion of female adult mosquitoes from the total population of early and late instar larvae (ranging between 0 and 16000) was used to compute the deterministic growth rate. This proportion was acquired by running the main text's Equations 1 and 2 to equilibrium, resulting in a population change from the initial condition to ~ 800 adults. Since pupae are not density-dependent, their abundance is not required in the simulation. Adult mosquitoes are also not density-dependent, but their abundance is essential for determining the Allee effect.

**Adult mosquitoes' population growth rate**

The per capita population growth rate estimates from the Leslie matrix increased with the size of the mosquito population to a maximum value, after which it declined as population size increased further. When the population was less than 200 mosquitoes, the growth rate remained below 1, reflecting the impact of the Allee effect below this population size. However, when the population increased between 200 and 600, the growth rate increased above 1, and as the population grew beyond 600, the growth rate decreased (Figure 5a). From an average of 100 simulations, at low population sizes, the growth rate was high, with values up to almost 2 (in part because the population will not be at a stable stage equilibrium), but declines sharply as the population size increases and eventually stabilises at around 1.0 for population sizes above 300 (Figure 5b).

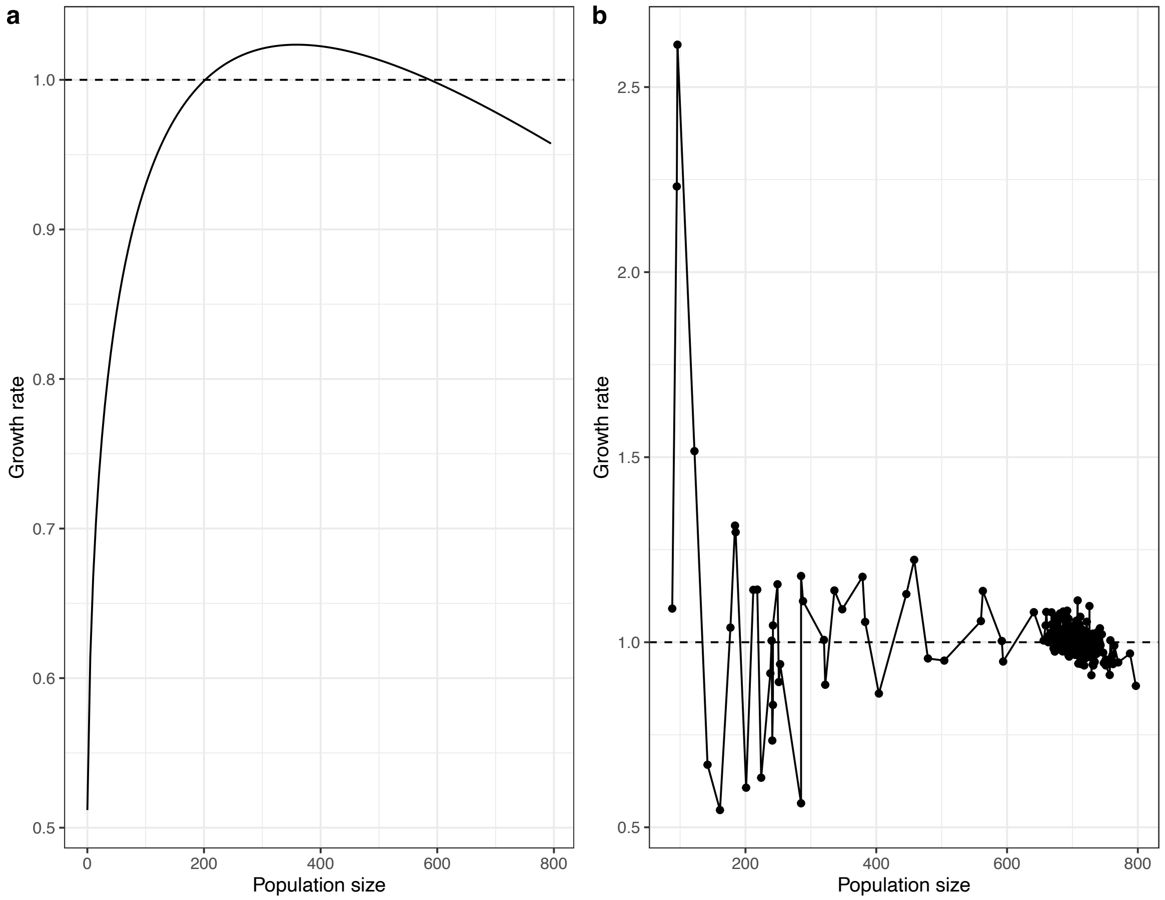


Figure 5: Population growth as estimated through (a) a Leslie matrix method and (b) direct calculations from the simulation model (i.e., an average across 100 simulations all initialised at 800 female adult mosquitoes). The dashed line indicates the threshold of the population growth rate, where the population declines when the rate is below this point.

**Trade-offs between negative density dependence and the Allee effects on the probability of extinction**

Since the real values of negative density-dependence and the Allee effect are unknown, negative density-dependence and the Allee effect sizes were varied from 100% reduction to 100% increase on their values set in the main text Table 1 (i.e., DD2 and AE2) and the probability of extinction was computed. A heatmap was used to show the probability of extinction across wards for each percentage change in both negative density-dependence and the Allee effect. For illustration, the simulation was repeated under three intervention regimes as described in the previous subsection, i.e., (a) without an intervention, (b) single short application of intervention, (c) double short applications of intervention and (c) sustained intervention (Table 1).

As negative density-dependence and the Allee effect increased, the probability of extinction also increased, but the Allee effect seemed to accelerate population extinction (Figure 6). While negative density-dependence in the absence of the Allee effect cannot lead the population to extinction, the Allee effect without density dependence can drive the mosquito population to extinction, especially with a sustained intervention (Figure 6). In the absence of density-dependence and the Allee effect (i.e., mean values 5.5e-05 and 360, respectively, reduced by 100%), the probability of extinction was 0 with and without larvicidal intervention (Figure 6a-d). Without intervention, the population could not go extinct unless density dependence and the Allee effect are both increased by at least 50% (Figure 6a). With a single short application of intervention, only a 30% increase in density dependence while keeping the Allee effect constant could drive the population to extinction with a probability of extinction ranging between 0.75 and 1.0 (Figure 6b). Similarly, with a double short application of intervention, only a 10% increase in negative density dependence while keeping the Allee effect constant could drive the population to extinction with a probability of extinction equal to 1 (Figure 6d). In Figure 5d, the mosquito population declined to extinction with a probability of extinction starting from 0.75 when both negative density-dependence and the Allee effect (i.e., 5.5e-05 and 360, respectively) increased by at least 5% with a sustained intervention. However, with nearly no density dependence, population size slowly declined to extinction if and only if the Allee effect was increased by 50% in the presence of larvicidal intervention (Figure 6c, d). Moreover, if density dependence is increased by at least 50%, population size declines to extinction even before reaching a 50% increase in the Allee effect, given there is an intervention

(Figure 6b-d). Furthermore, when density dependence and the Allee effect were both increased by at least 50%, their combination drove the population to extinction (Figure 6b-d). However, the presence of a sustained intervention accelerated extinction and made the combination of negative density-dependence and the Allee effect a threat to mosquito populations (Figure 6d).

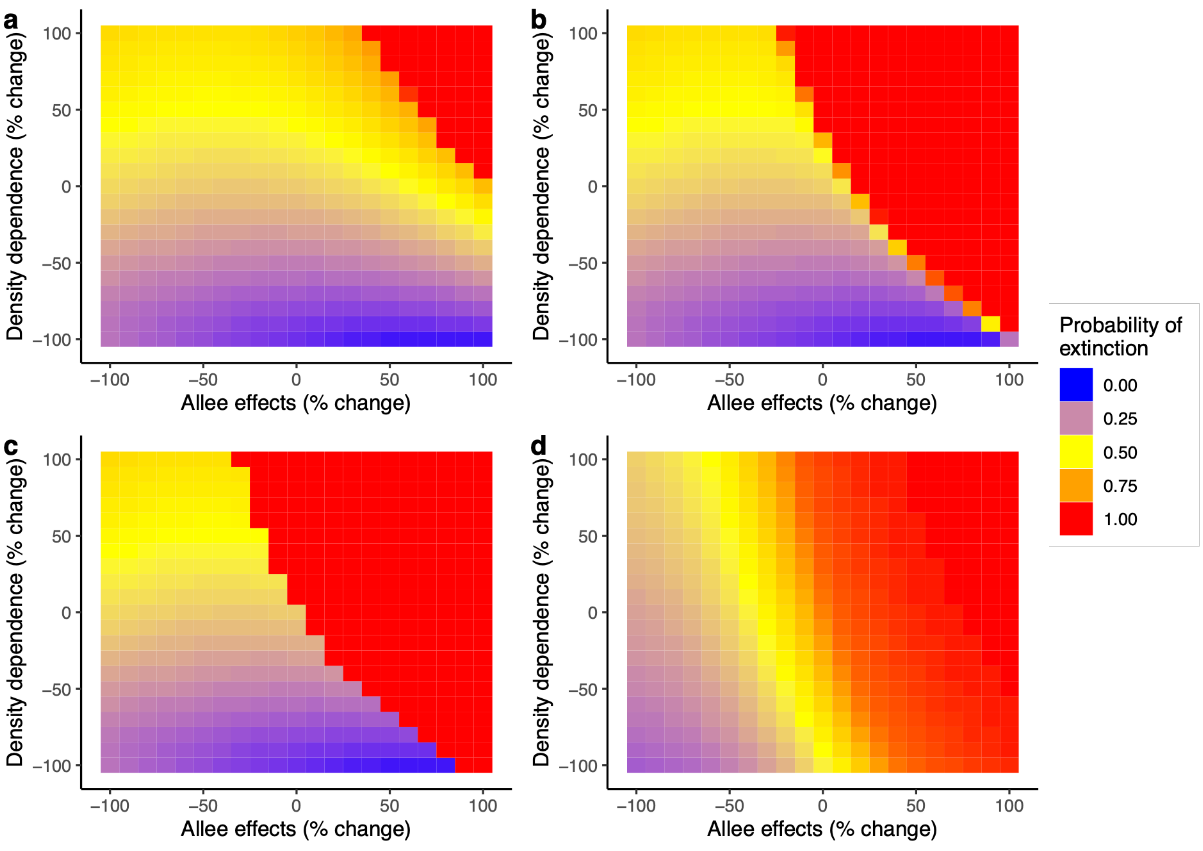


Figure 6: Heat maps showing the probability of extinction for each percentage increase or decrease in density-dependence and Allee effect mean values, 5.5e-05 and 360, respectively; (a) without intervention, (b) with a single short application of intervention, (c) double short applications of intervention and (d) sustained intervention.

**Replication of the results using higher initial population sizes relative to DD and AE values**

A. Role of DD and AE in regulating mosquito populations

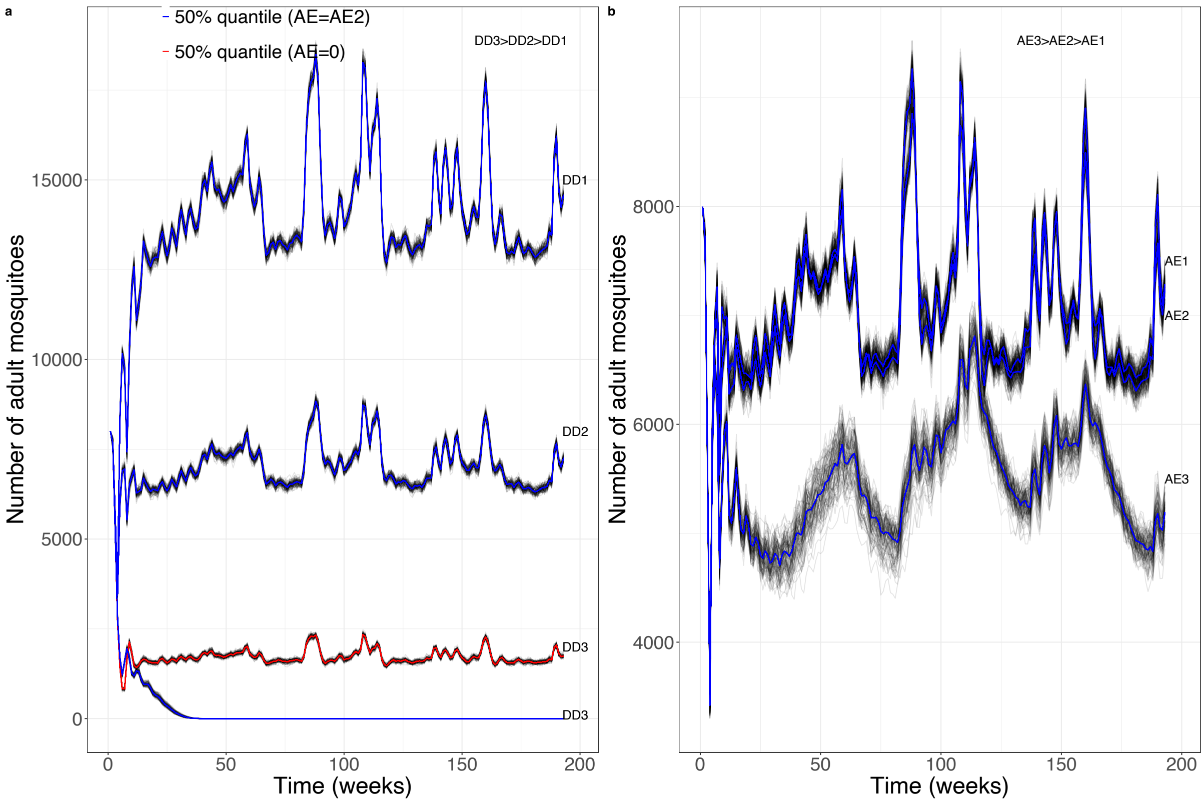


Figure 7: The role of DD and AE in regulating mosquito populations (without a larvicidal intervention) with higher population sizes relative to DD and AE values. (a) Three DD levels, DD1=2.75e-06, DD2=5.5e-06 and DD3=2.2e-05, were used, keeping AE constant at 3600 (AE2), and (b) Three AE levels, AE1=1800, AE2=3600 and AE3=7200, were used, keeping DD constant at DD2=5.5e-06. The red colour in (a) is the 50% quantile, which corresponds to DD3 when AE=0, and the blue colour in (a) and (b) is the 50% quantile when AE=AE2. (c) Percentage of population change (relative to the initial size of 8000 mosquitoes) after 193 weeks, with AE2 and DD2 varied across a range of values from 90% reduction to 100% or 300% increase. The percentage of population change was averaged over 100 simulations

### B. Impacts of negative density-dependence and Allee effects on sustained and short-term interventions

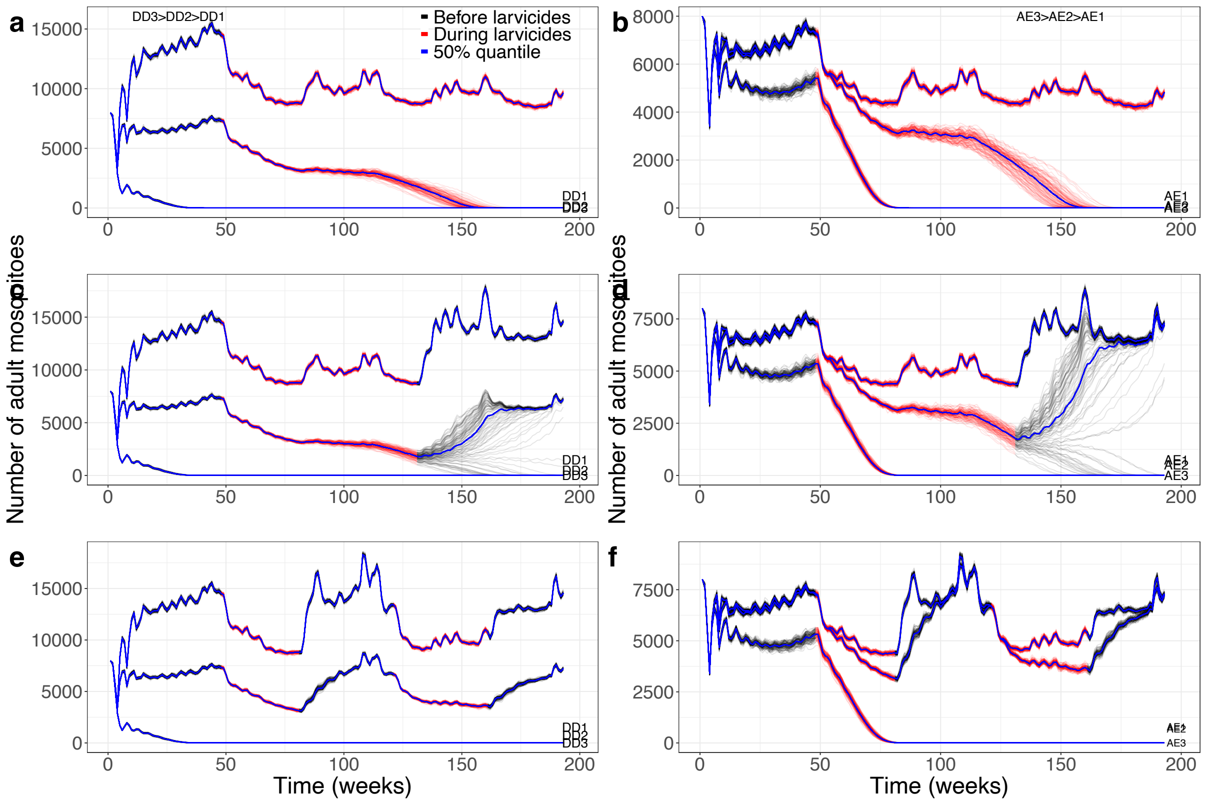


Figure 8: Mosquito populations regulated by (a, c, e) DD at levels 2.75e-06, 5.5e-06 and 2.2e-05, setting the AE (C) constant to AE2=3600 and (b, d, f) AE at levels 1800, 3600 and 7200, keeping DD constant at DD2=5.5e-06. Larvicidal treatment was applied in three regimes: (a, b) sustained larvicidal application, (c, d) a single short application and (e, f) two short applications.

### C. Trade-offs between negative density-dependence and the Allee effect in the mosquito population regulation

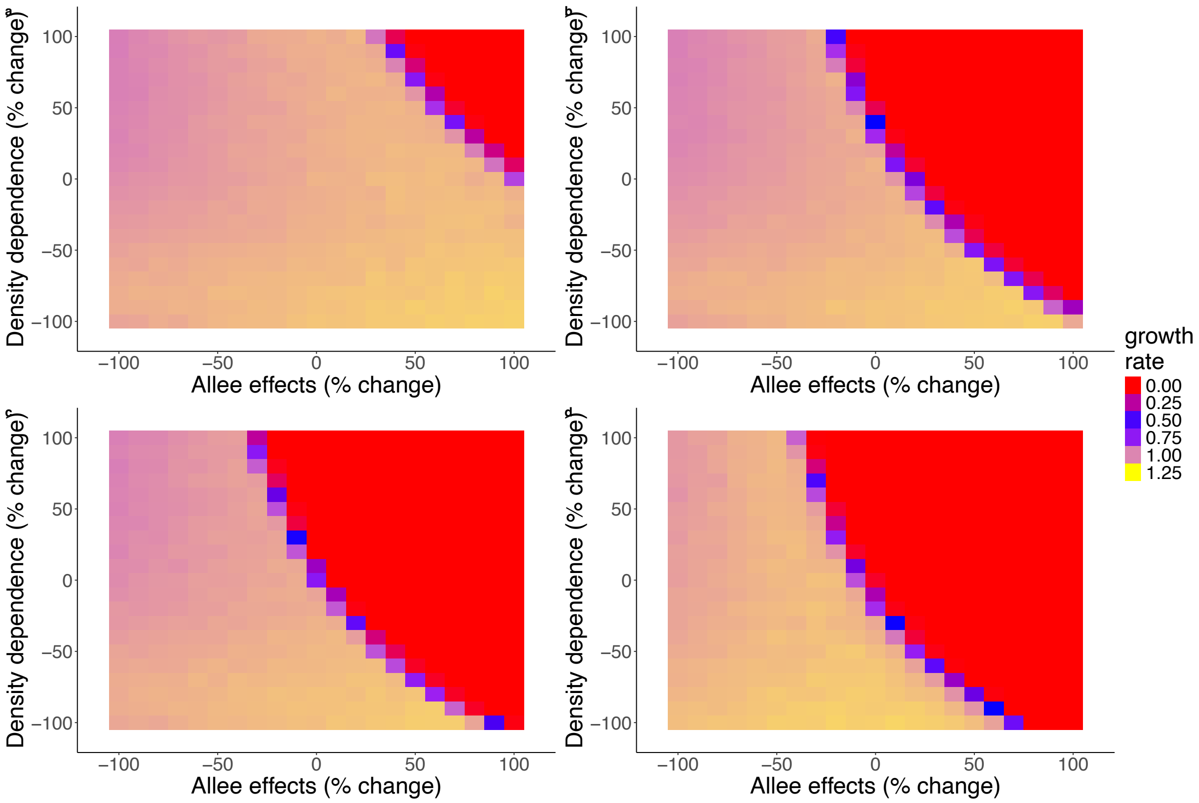


Figure 9: Heat maps showing the population growth rates averaged over 100 simulations for each percentage increase or decrease in DD and AE mean values, 5.5e-06 and 3600, respectively; (a) without intervention, (b) with a single short application of intervention, (c) two short applications of intervention and (d) with sustained intervention.

**An R script implementing the simulation model functions presented in the Results section**

```
#Intialization of time and number of wards
tmax=193 #Maximum time (in week)
Nwards=15 #Number of compartments, i.e, wards in our case

#Read rainfall and temperature data
rainfall = read.csv("weekly_rainfall.csv") #Read rainfall data
temperature = read.csv("weekly_Temperature.csv") #Read temperature data
R = as.data.frame(rainfall[,2]-56, drop=TRUE) #Write rainfall as a data frame
Te = as.data.frame(temperature[,2]-27, drop=TRUE) #Write rainfall as a data frame

#State vectors
Na<-matrix(0, Nwards, tmax) # Adult state vectors
Nl1<-matrix(0, Nwards, tmax) # Early instars state vectors
```

```
Nl2<-matrix(0, Nwards, tmax) # Late instars state vectors
Np<-matrix(0, Nwards, tmax) # Pupae state vectors

#Growth rate
Gr <- matrix(0,Nwards, tmax)

#Observation process
#Create an empty matrix with Nas in datNaPois for the Poisson process
datNaPois <- matrix(NA,Nwards, tmax)
datNaPois[1:Nwards]<-800

#Create empty matrix with Nas in datNaNB for the negative binomial process
datNaNB <- matrix(NA,Nwards, tmax)
datNaNB[1:Nwards]<-800

#Define initial values
Nl1[1:Nwards]<-1000      #Initialisation with earrly instars
Nl2[1:Nwards]<-1000     #Initialisation with late instars
Np[1:Nwards]<-900      #Initialisation with pupae
Na[1:Nwards]<-800      #Initialisation with female adults

#Definition and duration of experimental treatments
 LA<-matrix(0, Nwards, tmax)
 LA[1:3, 48:tmax]<-0 #First phase of larviciding
 LA[4:9,108:tmax]<-0 #Second phase of larviciding
 LA[10:15,155:tmax]<-0 #Third phase of larviciding

#Define a simulation function
simulation_function <- function(
  #Model parameters
  beta0, #Logit of baseline larval survival
  beta1, #Density dependence on larval survival
  beta2, #Interaction between larvae density and rainfall
  beta3, #constant defining sensitivity to rainfall in early instar larval survival
  beta4, #Effect of larvicides on early instar larval survival
  beta5, #Sensitivity of temperature to early instar larval survival
  lambda0, #Logit of baseline pupa survival
  omega0, #Log per capita fecundity
  omega1,#Constant defining the sensitivity of temperature to fecundity rate
  alpha0, #Logit of baseline adult survival
  he, #Hatching rate
  C, #Population size that scales the Allee effect
  stochastic # Simulates stochastic when TRUE and deterministic when FALSE
){
  require("abind")

#Define a loop for survivals and fecundity for total larvae, pupae and adults
  for(t in 2:tmax)
  {
   #Survival rates of early and late instars, pupae and adults
```

```
   sl1<-plogis(beta0-beta1*(1-beta2*R[t-1,])*(Nl1[,t-1] + Nl2[,t-1])+beta3*R[t-1,]-beta
4*LA[,t-1]+beta5*Te[t,])
   sl2<-plogis(beta0-beta1*(1-beta2*R[t-1,])*(Nl1[,t-1] + Nl2[,t-1])+beta3*R[t-1,]-beta
4*LA[,t-1]+beta5*Te[t,])
   sp<-plogis(lambda0)
   sa<-plogis(alpha0)

#Define survivors
   #Late instar larvae: Early to late instars
   Nl2[,t]<-
     if (stochastic) {
       rbinom(Nwards,Nl1[,t-1],sl1)
     } else{
       Nl1[,t-1]*sl1
     }

   #Pupae: Late instars to pupae
   Np[,t]<-
     if (stochastic) {
       rbinom(Nwards,Nl2[,t-1],sl2)
     } else{
       Nl2[,t-1]*sl2
     }

   #Total adults: from survived adults plus emerged from pupae
   Na[,t]<-
     if (stochastic) {
       rbinom(Nwards,Na[,t-1],sa)+rbinom(Nwards,Np[,t-1],sp)
     } else{
       Na[,t-1]*sa + Np[,t-1]*sp
     }

   #Fecundity rate: total number of eggs laid
   b<-exp(omega0+omega1*Te[t,])*Na[,t-1]*(Na[,t-1]/(C+Na[,t-1]))

   #Early instars: Eggs hatched to early instar larvae
   Nl1[,t]<-
     if (stochastic) {
       rpois(Nwards, 0.5*he*b)
     } else{
       0.5*he*b
     }

   #Define growth rate
   Gr[,t-1] <- Na[,t]/(Na[,t-1]+1)

  } # End of time t loop
  abind(LA=LA, Nl1=Nl1, Nl2=Nl2, Np=Np, Na=Na, Gr=Gr, along = 3)
} #End of simulation function
```

**Conversion of original parameter values to values per week**

One common challenge in calculating parameter values is the conversion of their observed or original values to a standardised time base, e.g., values per week. This can be done as follows:

$$Parameter\ value\ per\ week = (original\ biological\ value)^{\frac{7\ days}{t_0\ days}}$$

where $t_0$ is the original or observed time interval, such as one, two or three days.
